# A computational account of how positive performance bias supports cognitive effort

**DOI:** 10.64898/2026.05.13.725021

**Authors:** Kazuma Mori, Makiko Yamada

## Abstract

The willingness to exert cognitive effort is essential but is constrained by the subjective cost of effort. Although effortful tasks are often avoided, positive bias about one’s own performance may help sustain engagement with cognitive demands. Here, participants completed an effort-based decision-making task and reported trial-by-trial predictions of their own performance, allowing us to quantify performance prediction error (PPE) as the discrepancy between subjective and objective accuracy. The results showed that PPE was predominantly positive and increased with effort level, indicating greater overestimation under higher cognitive demands. Using a computational model, we show that choices were best explained by a learning model in which rewarded trials accompanied by positive PPE decreased subsequent sensitivity to effort. A confidence-based control model did not provide a better account of choices, suggesting that this effect was better captured by positive performance bias than by confidence alone. Our findings provide a computational account of how biased self-evaluation may attenuate the subjective cost of cognitive effort and extend the positive bias literature to the task need for cognitive effort.

## Introduction

In everyday life, people often face situations in which they must decide whether a potential outcome is worth the cognitive effort required to obtain it. Cognitive effort has been widely regarded as subjectively costly, and people tend to avoid cognitively demanding options when the expected benefits are held constant (Kool & Botvinick, 2018; Andrew Westbrook & Braver, 2015). This tendency has been formalized within effort-discounting frameworks, which describe how the subjective value of an outcome decreases as the effort required to obtain it increases. The effort-discounting paradigm has become an increasingly influential approach for quantifying how effort requirements shape choice behavior in humans (Escobar & Mitchell, 2025).

Importantly, effort sensitivity is not necessarily fixed. Rather, the effort-discounting parameter can fluctuate during task engagement as a function of recent experience (Sayalı & Badre, 2021). One prominent factor is fatigue. Computational studies have shown that momentary fatigue varies from trial to trial and predicts subsequent effort-based choices, suggesting that effortful exertion can increase the subjective cost of future action (Matthews et al., 2023; Müller et al., 2021). These findings indicate that effort-based decisions are governed not only by static effort and reward values, but also by dynamic learning processes that update effort sensitivity over time. In the present context, we focus on within-task fluctuations in effort sensitivity, rather than on stable differences in effort discounting across social contexts or task domains, such as decisions involving effort for oneself versus others (Lockwood et al., 2017, 2022).

Yet an important question remains unresolved: under what conditions do people seek, rather than avoid, effortful tasks? The effort paradox theory proposes that effort is not merely costly; it can also add value to outcomes and may itself become desirable in some contexts (Inzlicht et al., 2018). Consistent with this view, empirical studies have shown that exerting effort can enhance subsequent reward valuation (Ma et al., 2014) and that rewarding cognitive effort can increase later preference for more demanding tasks (Clay et al., 2022). These findings suggest that motivation to expend cognitive effort may be supported by psychological factors that counteract effort avoidance. Among such factors, positive bias in self-evaluation may be particularly important. Human self-evaluation is often positively biased: people tend to hold overly favorable views of themselves, their control over events, and their future (Sharot, 2011). Classic work on positive illusions further proposed that such biases are common in normal cognition and may contribute to psychological well-being and adaptive functioning (Taylor & Brown, 1988, 2004).

Positive bias may be particularly relevant to effortful behavior because persistence in difficult tasks often requires maintaining favorable beliefs about one’s own competence despite imperfect performance or repeated failures. If individuals believe that they performed better than they did, this biased self-evaluation may sustain the expectation that future effort will be successful. Conversely, depressive realism accounts suggest that individuals with depressive symptoms may show reduced positive bias or more realistic judgments under some conditions (Moore & Fresco, 2012). Relatedly, individuals with major depressive disorder have been reported to show greater effort discounting for rewards and reduced willingness to exert cognitive effort compared with healthy participants (Ang et al., 2023). Taken together, these findings raise the possibility that positive bias has an adaptive motivational function: overestimating one’s own performance may reduce the perceived cost of future effort, thereby promoting continued engagement with cognitively demanding tasks.

Despite this possibility, the role of positive bias in effort-based decision-making remains poorly understood. Whereas previous computational accounts have focused on how fatigue, effort exertion, or negative outcomes increase effort avoidance (Matthews et al., 2023; Müller et al., 2021), little is known about whether positively biased self-evaluation can exert the opposite effect by decreasing effort sensitivity and increasing subsequent willingness to work. This question can be addressed by measuring self-evaluation on a trial-by-trial basis and linking it to computational parameters that govern effort-based choice.

In the present study, we aimed to clarify whether positive bias about one’s own performance can counteract effort avoidance and support engagement with cognitively demanding tasks. Participants completed a cognitive effort decision-making task combined with trial-by-trial performance predictions. On each trial, they chose between a fixed rest option and a task option varied with effort and reward levels. After each cognitive effort task trial, participants predicted their own performance before receiving reward feedback. We defined performance prediction error (PPE) as the difference between subjective and objective accuracy, with positive values indicating overestimation of performance. Drawing on previous work on positive bias and positive illusions (Sharot, 2011; Taylor & Brown, 1988), we predicted that healthy participants would frequently overestimate their performance, allowing us to quantify positive bias on each trial and examine its influence on subsequent effort-based choices.

Using this task, we tested the central hypothesis that positive PPE following task performance would predict greater engagement in subsequent effort-based choices, particularly when later effort demands were high. To evaluate this hypothesis computationally, we developed a learning-based effort-discounting model in which the effort-discounting parameter was updated after each trial based on the trial outcome and PPE. We hypothesized that task failure on trial *t* would increase effort sensitivity and promote subsequent effort avoidance, consistent with previous work (Matthews et al., 2023), whereas rewarded trials accompanied by positive PPE on trial *t* would decrease effort sensitivity and thereby increase the subjective value of effortful options on later trials.

## Methods

### Participants

We recruited 46 healthy young adults (18 female, 28 male; mean age = 26.1 years, SD = 8.2). The study was approved by the QST Research Ethics Committee. Eleven participants were excluded for failing to follow task instructions, leaving 35 participants for the main analyses. Written informed consent was obtained from all participants. Participants received a fixed compensation of 10,000 yen. No additional performance-contingent payment was provided, although previous work has shown that participants can perform better even in the absence of monetary incentives (Gneezy & Rustichini, 2000).

### Cognitive effort task

To assess cognitive effort, we used a rapid serial visual presentation (RSVP) task (Figure 1). Whereas previous studies manipulated effort by varying the number of attentional switches (Apps et al., 2015; Chong et al., 2017; Yantis et al., 2002), our adapted version manipulated the speed of the stimulus stream. By adjusting the RSVP stream speed on each trial, we varied the required level of cognitive effort: a stimulus duration of 490 ms corresponded to the lowest effort level, whereas 350 ms corresponded to the highest. Participants fixated centrally while arrays of letters were presented sequentially, with 40 stimulus presentations per trial over 14–19.6 s. In a preliminary experiment, we confirmed that task accuracy could be modulated by these effort settings. Because a central aim of the present study was to dissociate confidence from objective performance across easy and difficult conditions, we adopted this speed-based manipulation rather than the attentional-switch manipulation used in previous studies (Apps et al., 2015; Chong et al., 2017; Yantis et al., 2002). Importantly, the preliminary experiment also confirmed that participants were willing to perform the low-effort condition despite its longer trial duration (i.e., effort level 1 was 5.6 s longer than effort level 5). Thus, although rewards are known to be devalued by delays in receipt (Green et al., 1994), the present design minimized the possibility that choices between task and rest were driven primarily by temporal discounting.

**Figure 1.**
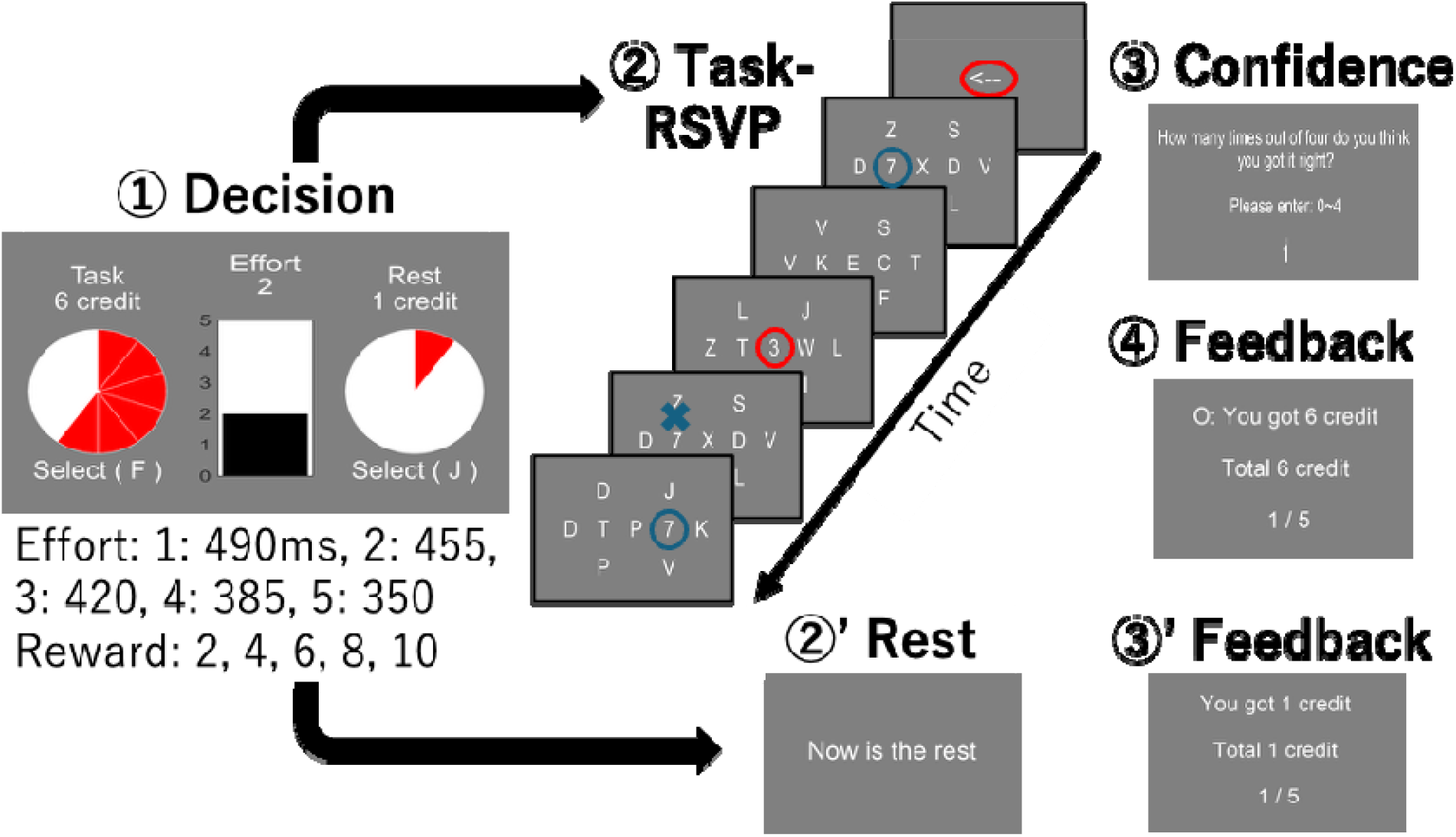
Cognitive effort-based decision-making task. Each trial began with a decision phase in which participants chose between a task option that varied in cognitive effort and reward magnitude and a fixed no-effort, low-reward rest option. The task option required participants to perform a rapid serial visual presentation (RSVP) task. During the RSVP task, participants were asked to respond to one of two lateral target letter streams when the target number “7” appeared. The currently attended target stream is indicated by the blue circle in the Task-RSVP panels. An arrowhead presented at the beginning of the RSVP trial indicated the initial target stream, and the appearance of a central “3” instructed participants to shift attention to the opposite target stream. Cognitive effort was manipulated by varying the presentation speed of the RSVP streams, with faster streams corresponding to higher effort levels. After completing the task, participants rated their confidence in their performance and then received feedback indicating whether they had earned the offered reward. If participants chose the rest option, they rested briefly and received the fixed low reward.

The main task required participants to fixate centrally while monitoring two target letter streams located to the left and right of fixation for the appearance of the target number “7”. Each target stream was surrounded by three distractor streams. Four targets were presented on each trial, separated by 3–12 streams, and participants pressed the space bar whenever they detected a target. Successfully responding while “7” was on the screen was counted as a correct response. At the beginning of each trial, a left- or right-pointing arrow was presented at fixation for 700 ms to indicate the target stream to which participants should initially attend. During the trial, participants were required to shift attention to the opposite target stream whenever the number “3” appeared at fixation. Four attentional switches occurred on each trial, also separated by 3–12 streams. After reporting their confidence in task performance (see Experimental procedure), participants received feedback indicating whether they had earned a point. To earn a point, participants had to detect at least one of the four targets. Rapid consecutive keypresses were disabled, and trials with more than six space-bar presses were not permitted.

### Stimuli and apparatus

Participants were seated approximately 65 cm from the 24-inch display. Three distractor streams flanked each peripheral stream, positioned above, below, and lateral to the target stream. The target streams were positioned approximately 3° to the left and right of the central array, with lateral distractor streams extending farther into the periphery. These distractor streams presented task-irrelevant stimuli to increase task difficulty. Stimuli consisted of randomly selected white letters presented in Arial font on a gray background, with a text height of 0.06. Previous work has shown that only the central and peripheral streams can be attended simultaneously (Yantis et al., 2002), requiring participants to shift attention to detect targets. Experiments were conducted on a PC running PsychoPy (www.psychopy.org), which was used for stimulus presentation and response collection.

### Experimental procedure

In the effort discounting task, participants made cost-benefit decisions (Figure 1) between a “rest” option requiring no effort and yielding a low reward (1 credit) and a “task” option requiring greater effort (levels 1–5) with the RSVP task and yielding a higher reward (2, 4, 6, 8, or 10 credits). The effort discounting task allowed us to quantify effort and reward sensitivity. Across trials, the effort and reward levels of the offer varied, whereas the rest option remained constant. Participants completed 75 trials, comprising three repetitions of each effort-reward combination. The task and rest options were presented centrally as black bars on a white background, with bar height indicating effort level and credits displayed in a pie chart on the left. Choices were made using two keys on a mini keyboard corresponding to the right-hand side of the screen. Because the preliminary experiment indicated that participants were sufficiently motivated without a performance bonus, no additional payment was provided based on task performance. The total number of trials, including practice, was limited to 80 to minimize fatigue-related influences on task/rest decisions (Müller et al., 2021). After each RSVP trial, participants reported their confidence (0–4) about their task performance by answering the question, “How many times out of four do you think you got it right?” Finally, participants received feedback indicating whether they had earned credits. To prevent explicit score feedback from making subsequent predictions trivial, feedback indicated only whether credits had been earned, not the number of correct responses.

All participants first completed a brief practice session, followed by a test session. The practice session consisted of five trials, with the identical procedure as the test session. The practice began with a trial containing an effort level 1 with the slowest RSVP speed, and the effort level increased by 1 on each subsequent trial.

### Performance prediction error (PPE)

In the present study, we operationalized positive performance bias using performance prediction error (PPE), defined as the difference between subjective accuracy and objective accuracy in the RSVP task. Subjective accuracy was indexed by participants’ confidence ratings regarding the number of correctly detected targets, whereas objective accuracy was defined by actual task performance. Positive PPE values indicate that participants overestimated their performance relative to their objective accuracy. Because PPE may also reflect factors such as task difficulty, uncertainty, or individual differences in confidence scale use, we treated PPE as an operational index of positive performance bias rather than as a direct measure of positive bias itself.

### Statistical test

We used R (version 4.5.3) with RStudio (version 2026.04.0+526) for analysis. We analyzed behavioral choice data and PPE scores with general linear mixed-effects models (lme4 package, 2.0-1). We used logistic mixed-effects models to analyze participants’ decisions to select a task or rest, whereas we used linear mixed-effects models to analyze PPE scores, calculated as the difference between confidence and behavioral accuracy on the RSVP task. For all GLMMs, subject-specific intercepts are set as a random effect.

### Computational modelling

We tested several computational models employing a hierarchical Bayesian approach that simultaneously estimated decision processes at the group and individual levels to more accurately capture behavioral variation arising from the discounting of rewards by effort. Each of these models assumes that the probability of choosing an option is a function of its subjective value, which depends on the option’s reward and effort.

To characterize how positive bias modulates the motivation to exert cognitive effort, we mainly tested learning-based computational models. These models were designed to explain participants’ decisions (task or rest) by allowing the subjective value of the task option to vary as a function of reward, effort, trial outcome, and PPE. We investigated the hypothesis that the motivation represented by the effort-discounting parameter would change over the course of the task, influenced by both trial outcome and PPE.

### Baseline model selection

Before examining learning-based models, we first fit standard effort-discounting models to estimate choice parameters in the absence of PPE and trial outcomes. These models have been widely used to characterize how individuals trade off rewards against effort (Chong et al., 2017; Cutler et al., 2025; Lockwood et al., 2022). They assume that the subjective value of an option scales with the offered reward and is discounted as a function of the required effort. The form of the discounting function determines how effort shapes choice behavior. A linear model assumes a constant discount rate as effort increases. By contrast, hyperbolic and exponential models imply greater sensitivity to changes at lower effort levels than at higher levels, whereas a parabolic model implies the reverse. These functional forms have been widely adopted in previous studies of cognitive effort discounting (Chong et al., 2017; Cutler et al., 2025; Lockwood et al., 2022).

We compared three models using linear, parabolic, and hyperbolic effort-discounting functions. For each model, we fit participants’ task/rest choices in the cognitive effort task. Hierarchical Bayesian model comparison indicated that the parabolic discounting function provided the best fit to the data:

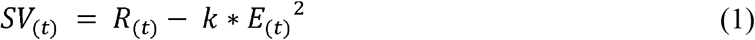

where Subjective value (*SV*) was estimated as the reward (*R*)offered on a given trial, discounted by the associated effort (*E*)(Model 1). The effort-discounting parameter *k* governed the extent to which rewards were devalued by effort. Because subjective values decrease as *k* increases, this parameter indexes the degree to which each participant discounted rewards with respect to effort. Note that the model hypothesized that *k* is static and does not fluctuate during the cognitive effort task.

To transform subjective values into choice probabilities, *P*_*(Task)*_, we used a softmax function:

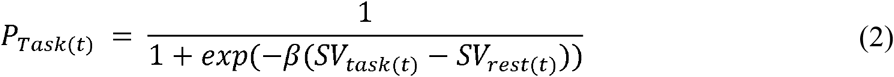

Here, *β* represents the inverse temperature parameter of the softmax function. The parameter captured the stochasticity of participants’ choices: higher β values indicate more consistent selection of the option with the higher subjective value, whereas β values close to zero indicate more random choice behavior.

### Learning-based modeling through PPE scores and trial-by-trial outcomes

To estimate fluctuations in motivation in effort-based decisions, we adapted Eq. 3 to allow rewards to be discounted by effort on a trial-by-trial basis:

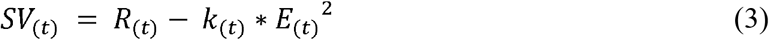

The key difference between Eq. 1 and Eq. 3 is the time-varying discount factor *k*_(*t*)_. Thus, the trade-off between reward and effort changes across trials as a function of *k*_(*t*)_. The evolution of this discount factor defined how each model explained trial-by-trial choices. Initial k values were constrained to the range 0 to 3, following previous work (Todorova et al., 2025).

Using trial-by-trial *k*_(*t*)_ values, we first tested the model from a previous study (Matthews et al., 2023), which assumes that an individual’s effort-discounting parameter increases after unrewarded trials. This model proposes that task motivation depends on whether participants successfully performed the task:

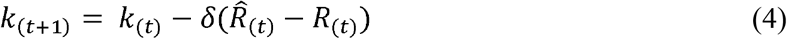

where *δ* denotes the learning rate (0 < *δ* ≤ 1), and *R*_(*t*)_ and 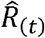 denote the chosen and obtained rewards on the previous trial, respectively. 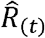 is equal to the chosen reward when the participant succeeds and equal to 0 following an error. We therefore implemented an error-driven, effort-aversive learning rule in which *k*_(*t*)_ increases after trials in which the participant fails (Model 2). Under this formulation, unsuccessful outcomes progressively increase the model’s sensitivity to effort.

We next evaluated a novel model in which the effort-discounting parameter was assumed to decrease after rewarded trials accompanied by PPE. Under this account, task motivation is selectively shaped by the magnitude of PPE, but only when participants successfully complete the task:

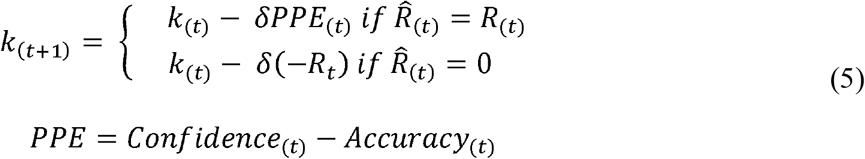

where PPE represents prediction error in task performance. In this equation (Model 3), we additionally incorporated a PPE-based effort-seeking learning term, such that *k*_(*t*)_ decreases after trials in which participants overestimate their performance. This term was applied only when PPE was positive, so that overestimation of performance selectively reduced effort sensitivity. Under this formulation, positive bias about task performance progressively reduces the model’s sensitivity to effort. As a control model, we also evaluated a confidence-based learning model in which the PPE term was replaced with trial-by-trial confidence ratings (Model 4). This model tested whether the effort-seeking effect was attributable to subjective confidence itself rather than to positive performance bias.

Note that we set the same free parameter for both effort-aversive and seeking learning, based on the assumption of shared learning sensitivity, as in a basic reinforcement learning model (Sutton & Barto, 1998). In practical terms, since the number of trials is limited in this study, this setting is effective for improving parameter identifiability. We also found that the different learning parameter models, each for effort-aversive and seeking, did not provide a good fit.

### Model fitting and selection

Model parameters (*k, β, δ*) were estimated separately for each participant using hierarchical Bayesian modeling. Model estimation was performed in R using the rstan package (Stan Development Team, 2026), following best practices implemented in hBayesDM (Ahn et al., 2017). We ran four independent Markov chain Monte Carlo (MCMC) chains, each with 2,000 iterations, discarding the first 1,000 iterations of each chain as warm-up. The remaining 1,000 iterations per chain were retained, yielding 4,000 posterior samples. All models reported in the main manuscript showed satisfactory convergence, with R □< 1.01. For model selection, we used the leave-one-out information criterion (LOOIC) (Todorova et al., 2025; Vehtari et al., 2024), which evaluates predictive accuracy while penalizing model complexity to avoid overfitting. Lower LOOIC values indicate better predictive performance.

### Model validation and parameter recovery

We performed posterior predictive checks and a parameter recovery analysis to validate the models, using data generated by the winning model and simulated participants. This synthetic dataset was created by randomly sampling individual parameter values from the posterior group-level ground-truth estimates derived from the winning model. We simulated choice probabilities to assess whether the generated data replicated key patterns observed in participant behavior as a function of the effort-reward combination. We also assessed subject-level parameter recovery by computing the correlation between the true and simulated subject-level parameter estimates.

## Results

The present study examined how positive bias arising from performance prediction error (PPE), defined as the difference between subjective accuracy (confidence) and objective accuracy (behavioral performance), fluctuates from trial to trial and influences the subjective value of exerting cognitive effort to obtain rewards. We used an effort discounting decision-making task in which participants chose whether to exert cognitive effort to obtain a reward (Figure 1). In the main task, participants were not allowed to take breaks; instead, on each trial, they chose either to perform the task or to rest. Tasks offer varied rewards in both reward magnitude (2, 4, 6, 8, or 10 credits) and the level of cognitive effort required (1, 2, 3, 4, or 5). Declining the task resulted in a no-effort rest option that yielded a small reward of 1 credit.

Cognitive effort was manipulated using the RSVP task. On each trial, participants were presented with task offers that varied in effort level, corresponding to higher or lower mental demand. Mental demand was manipulated by the RSVP stream’s speed. To assess trial-by-trial changes in positive bias, participants answered their confidence about accuracy on the RSVP task on a scale from 0 to 4 after each trial and before receiving reward feedback.

### Manipulation check

To analyze task/rest choices in the effort task, we used a logistic mixed-effects model with effort and reward levels as fixed effects. Participants were less likely to choose the effortful option as effort increased (OR = 0.27, 95% CI [0.20, 0.36], *p* < 0.001) and more likely to choose it as reward increased (OR = 2.35, 95% CI [1.78, 3.09], *p* < 0.001). The effort × reward interaction was also significant (OR = 0.92, 95% CI [0.87, 0.98], *p* = 0.01), indicating that rewards became less effective incentives as effort increased (Figure 2). These results confirmed that participants performed the task with recognition of the effort-reward cost-benefit as similar as previous studies (Apps et al., 2015; Chong et al., 2017; Cutler et al., 2025; Lockwood et al., 2022).

**Figure 2.**
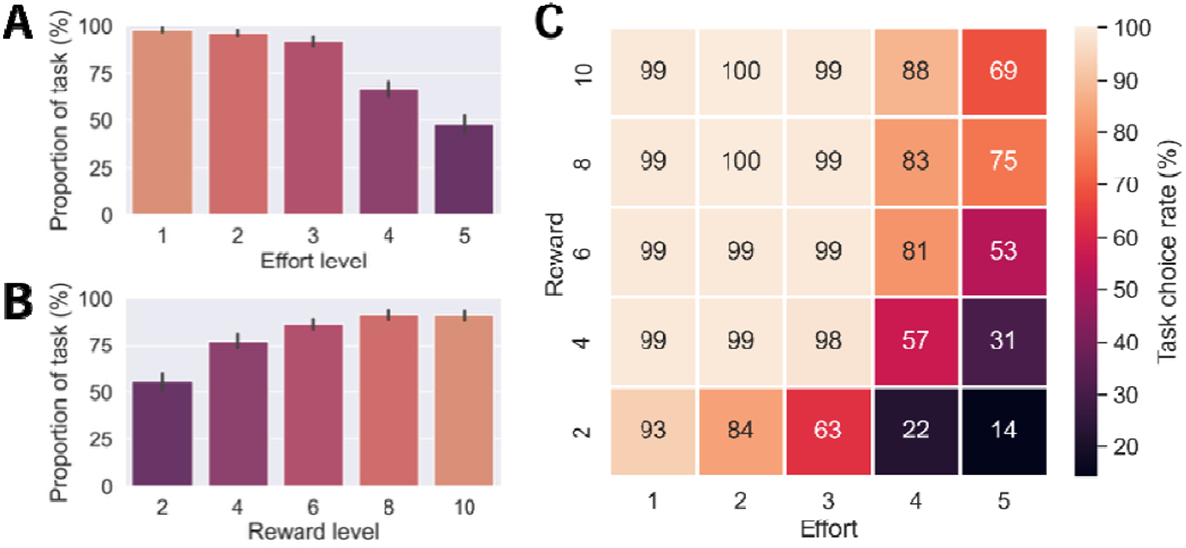
Effort and reward modulate decisions to engage in the cognitive effort task. Mean proportion of accepted task offers as a function of (A) effort level, (B) reward level, and (C) the combination of effort and reward level. Error bar indicates a 95% confidence interval.

### PPE showed a robust positive bias and increased in proportion to the effort level

Next, we examined whether PPE reflected a positive bias in the cognitive effort task. The distribution of PPE scores showed that 60% were greater than 0, 30% were equal to 0, and 10% were less than 0 (Figure 3A). This distribution, which was positively skewed (skewness = 0.26, z = 4.71, *p* < 0.001), indicates that participants often overestimated their own performance.

**Figure 3.**
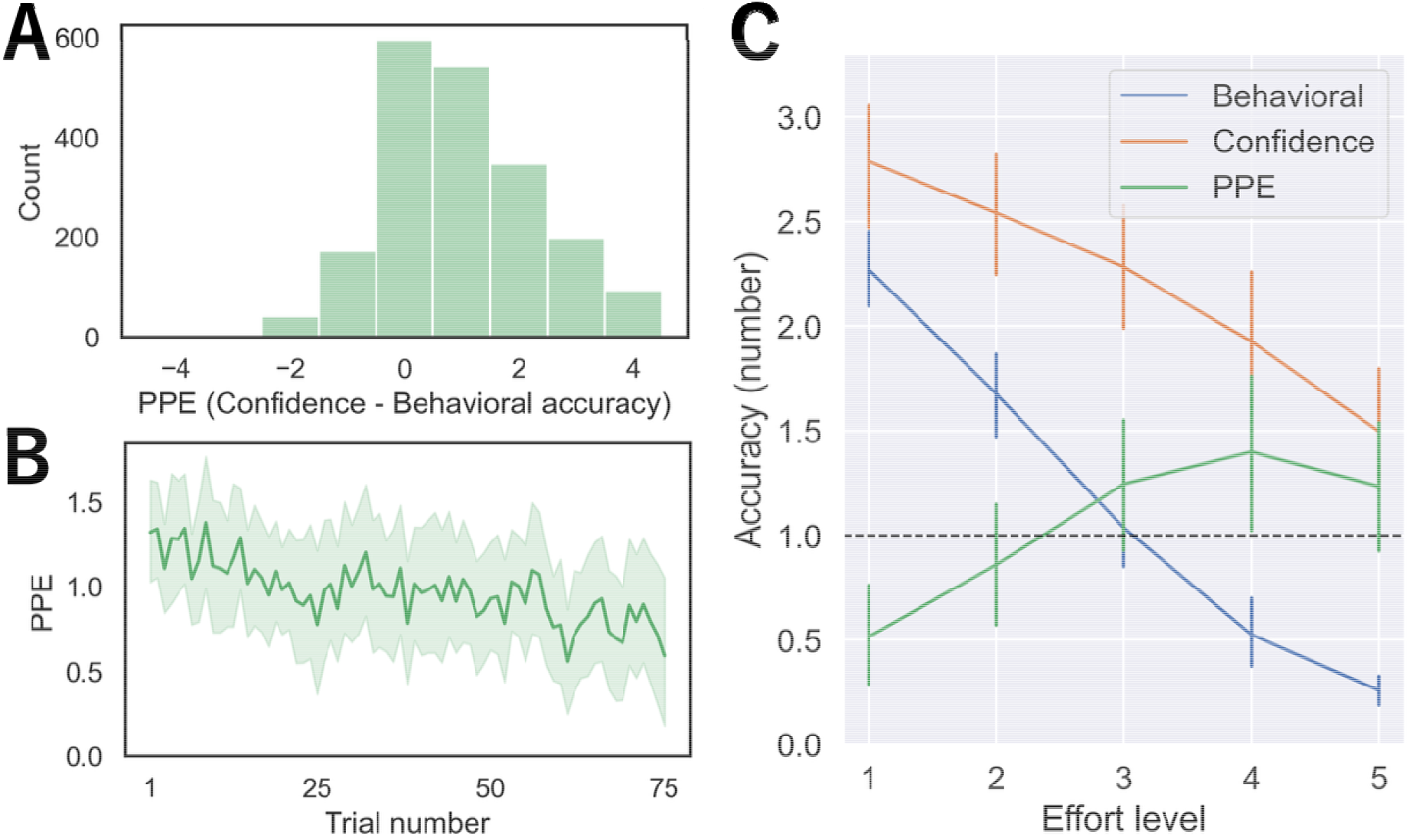
Performance prediction error reflects a persistent positive bias that increases with cognitive effort. (A) The distribution of PPE (defined as subjective accuracy minus objective accuracy). Positive values indicate overestimation of task performance. (B) Trial-by-trial change in PPE. PPE decreased over time but remained significantly above zero at the final trial. (C) PPE as a function of effort level. PPE increased with effort, indicating that positive bias became stronger under more effortful conditions.

To show whether the positive bias remained in later trials, we then used a linear mixed-effects model to examine trial-by-trial changes in PPE, with trial number as a fixed effect. The time-series data were smoothed using a five-point moving average. PPE decreased significantly over the course of the task (*β*= -0.005, 95% CI [-0.007, -0.003], *p* < 0.001; Figure 3B), indicating that overestimation of task performance gradually declined with experience. Because PPE was obtained only when participants chose the task option, PPE was available for 26 participants who selected the task on the final trial. Among these participants, PPE remained significantly above zero at the final trial, *t*(25) = 2.55, p = 0.02, suggesting that positive bias was not fully corrected even after participants had accumulated substantial metacognitive experience with the task.

Furthermore, we tested whether PPE was modulated by effort and reward levels using a linear mixed-effects model. PPE increased significantly with effort (*β*= 0.27, 95% CI [0.17, 0.37], *p* < 0.001; Figure 3C), indicating that positive bias became stronger as cognitive effort increased. The main effect of reward (*p* = 0.14) and the effort × reward interaction (*p* = 0.50) was not significant.

### Computational modeling revealed the effort-seeking effect of PPE

We first examined whether choice data were best explained by a linear, parabolic, or hyperbolic effort-discounting function, with a fixed effort-discounting parameter across trials. The best-fitting baseline model was one in which effort discounted rewards according to a parabolic function (Model 1, LOOIC = 1357; Figure 4A), consistent with previous studies of cognitive effort discounting (Ang et al., 2023; Matthews et al., 2023). Fitting choices without learning rules allowed us to estimate each participant’s baseline effort aversion and choice stochasticity, which were then used in the subsequent learning-rule models.

**Figure 4.**
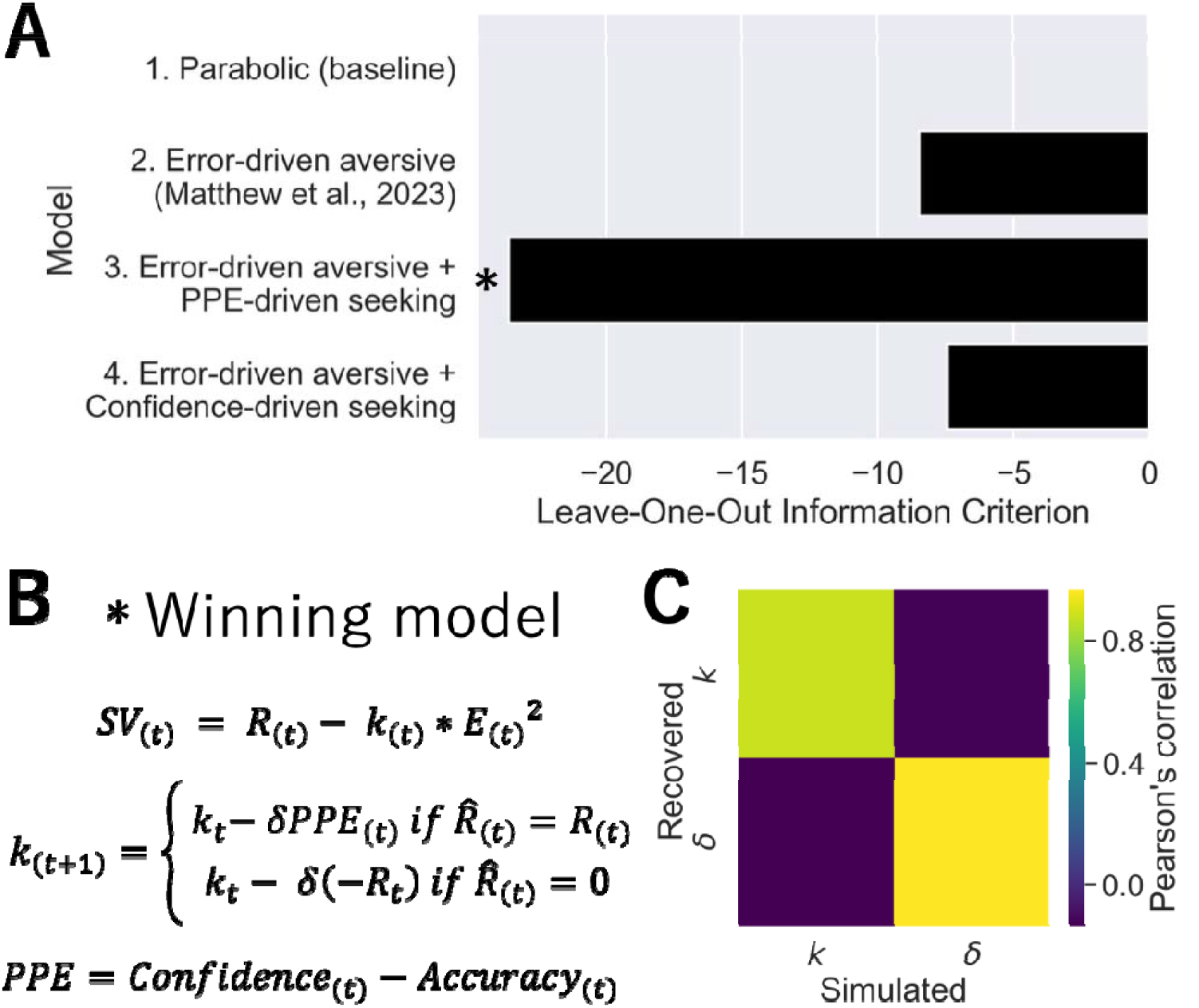
Computational modeling indicates that positive PPE may reduce subsequent effort sensitivity. (A) Computational models were compared to determine how effort discounting changed across trials as a function of task outcomes and positive bias. (B) Model comparison based on LOOIC showed that the winning model incorporated both error-driven effort avoidance and PPE-driven effort seeking. (C) Parameter recovery analysis for the winning model. The strong correlations between simulated and recovered parameters supported the robustness and identifiability of the model parameters.

We then fit computational models to PPE scores and effort-based decisions to examine whether they could account for fluctuations in choice behavior. We developed three trial-by-trial learning models (Model 2-4, Figure 4A) in which trial outcomes updated the effort-discounting parameter independently of effort level. Model 2 tested the hypothesis, based on previous work (9), that the effort-discounting parameter increases after failed trials, reflecting error-driven effort avoidance and reducing motivation to engage in the task on subsequent trials. Model 3 extended this framework by testing whether positive PPE on trial *t*, when the participant successfully obtained the reward, decreased the effort-discounting parameter on the next trial, *k*_*t+*1_. This PPE-driven update increased the subjective value of future task options and thereby the probability of choosing the task on subsequent trials. This PPE-driven effort-seeking mechanism increases the subjective value of future task options, thereby increasing the probability of choosing the task. Finally, Model 4 also tested a confidence-driven effort-seeking mechanism to clarify the importance of PPE.

Consistent with our predictions, choices were best explained by a delta-learning rule that captured the combined effects of errors and positive bias (Model 3, LOOIC = 1333; Figure 4A, B). For the winning model, the median point-biserial correlation between observed and predicted choices was 0.89 (SD = 0.07). This model outperformed the alternatives and showed satisfactory recovery of the main parameters (*k, r* = 0.88 and *δ, r* = 0.97; Figure 4C). Because Model 4 did not outperform Model 2, the effort-seeking effect was specific to PPE and was not explained by confidence alone.

## Discussion

The present study examined whether positive bias about one’s own performance is related to the willingness to engage in cognitively effortful behavior. We found that participants frequently overestimated their task performance, and this positive performance bias increased as cognitive effort demands increased. Computational modeling further suggested that choices were best explained by a learning model in which unsuccessful outcomes increased subsequent effort sensitivity, whereas rewarded trials accompanied by positive PPE decreased subsequent effort sensitivity. These findings suggest that positive performance bias may help sustain cognitive effort by attenuating the subjective cost of future effortful choices.

The present findings extend the literature on positive bias by showing that overestimation of one’s own performance emerges robustly during cognitively effortful behavior. PPE was predominantly positive and remained above zero even after repeated task experience, suggesting that participants did not fully recalibrate their self-evaluation despite receiving feedback throughout the task. This persistence is consistent with the view that positively biased self-evaluation is a common feature of normal cognition and may support adaptive functioning (Taylor & Brown, 1988, 2004; Sharot, 2011). Importantly, PPE increased with effort level, indicating that positive performance bias was not simply a stable individual tendency but varied systematically with task demand. This finding suggests that demanding cognitive contexts may amplify the discrepancy between subjective and objective performance, either because objective performance declines with higher effort or because individuals maintain favorable beliefs about their competence as tasks become more challenging. Thus, positive performance bias may be especially relevant in the very situations where effort avoidance would otherwise be expected.

These findings also contribute to the effort paradox literature by identifying positive performance bias as a potential psychological factor that may help sustain engagement with demanding tasks. Previous work has shown that effort can be costly, reducing the subjective value of rewards, but can also acquire value depending on context, learning, and subjective appraisal (Inzlicht et al., 2018; Ma et al., 2014; Clay et al., 2022). The present study suggests a complementary route by which effortful engagement may be maintained: individuals may continue to approach cognitively demanding tasks when they evaluate their own performance more favorably than it warrants based on objective accuracy. This does not imply that positive bias makes effort intrinsically rewarding. Rather, positive bias may attenuate the subjective impact of effort by sustaining the belief that future task performance will be successful. In this sense, positive performance bias may help bridge the apparent conflict between effort as a cost and effort as something people sometimes continue to pursue.

Although we did not measure neural activity, the present computational account is consistent with neural models of effort-based decision-making. Prior studies have implicated dorsal anterior cingulate and medial frontal regions in representing effort costs and guiding cognitive control allocation, while striatal and dopaminergic systems have been linked to the integration of reward benefits and effort costs (Chong et al., 2017; Hauser et al., 2017; Lopez-Gamundi et al., 2021; Westbrook et al., 2020). Our model suggests that positive performance bias may reduce subsequent effort sensitivity, potentially by altering value computations in cortico-striatal circuits. This possibility aligns with evidence that cognitive effort and reward anticipation share overlapping neural systems (Vassena et al., 2014), and that dynamic changes in motivation and fatigue can be modeled through trial-by-trial updates in effort valuation (Müller et al., 2021; Matthews et al., 2023). Future studies combining the present task with fMRI or EEG could examine whether PPE-related updates in effort sensitivity are encoded in medial frontal, insular, and striatal systems during subsequent choices.

The computational modeling results provide a mechanistic interpretation of this possibility. Consistent with previous accounts of effort avoidance and fatigue, failed trials were modeled as increasing subsequent effort sensitivity, thereby reducing the subjective value of future task options (Müller et al., 2021; Matthews et al., 2023). Critically, choices were best captured by a model in which rewarded trials, accompanied by positive PPE, decreased subsequent sensitivity to effort. This model-based result suggests that positive performance bias may operate as a countervailing update signal against effort avoidance: unsuccessful outcomes make effort more costly, whereas overestimation following successful performance makes future effort less costly. The confidence-based control model did not provide a better account of choices, indicating that the relevant signal was not confidence per se but the discrepancy between subjective and objective performance. Thus, the computational results support the idea that positive performance bias may influence effort-based decision-making by dynamically modulating the subjective cost of cognitive effort.

Although we did not measure neural activity, the present computational account is consistent with neural models of effort-based decision-making. Prior studies have implicated dorsal anterior cingulate and medial frontal regions in representing effort costs and guiding cognitive control allocation, while striatal and dopaminergic systems have been linked to the integration of reward benefits and effort costs (Chong et al., 2017; Hauser et al., 2017; Lopez-Gamundi et al., 2021; A. Westbrook et al., 2020). Our model suggests that positive performance bias may reduce subsequent effort sensitivity, potentially by altering value computations in cortico-striatal circuits. This possibility aligns with evidence that cognitive effort and reward anticipation share overlapping neural systems (Vassena et al., 2014), and that dynamic changes in motivation and fatigue can be modeled through trial-by-trial updates in effort valuation (Müller et al., 2021; Matthews et al., 2023). Future studies combining the present task with neuroimaging could examine whether PPE-related updates in effort sensitivity are encoded in medial frontal, insular, and striatal systems during subsequent choices.

Several limitations should be noted. First, the present study is correlational regarding the trial-by-trial relationship between PPE and subsequent choice. Although the computational model suggests that positive bias updates effort sensitivity, the design does not establish a causal effect of positive bias. Future studies could manipulate feedback, confidence, or perceived performance to test whether experimentally induced positive bias increases effortful choice.

Second, PPE was defined as the difference between self-reported confidence and objective accuracy. Although this operationalization directly captures overestimation, confidence ratings may also reflect task difficulty, response uncertainty, or individual differences in scale use. Future work should examine whether similar effects are obtained using more refined metacognitive measures, such as confidence calibration or metacognitive sensitivity. Third, the task involved healthy young adults and a specific RSVP-based manipulation of cognitive effort. It remains unclear whether the same mechanism generalizes to other types of cognitive effort, physical effort, older adults, or clinical populations. Fourth, although the winning computational model showed good predictive performance and satisfactory parameter recovery, the number of trials was limited, and the same learning-rate parameter was used for both effort-avoidance and effort-seeking updates to improve identifiability. Future studies with larger datasets could test more flexible models with separate learning rates for negative outcomes and positive bias.

In conclusion, the present study shows that positive bias in self-evaluation is not merely a metacognitive error but may play an adaptive role in effort-based decision-making. Participants frequently overestimated their performance, especially under higher cognitive effort, and choices were best explained by a model in which positive PPE attenuated subsequent effort sensitivity.

Computational modeling further indicated that rewarded trials accompanied by positive PPE reduced effort sensitivity, counteracting the effort-avoidance effect of task failure. These findings suggest that biased self-evaluation can dynamically regulate the subjective cost of cognitive effort and help sustain engagement with demanding tasks. More broadly, the results provide a computational account of how positive performance bias may support motivation by making effortful action feel more worthwhile.

## Acknowledgements

This study was supported by grants from the JST Moonshot R&D Grant (JPMJMS2295) and CREST Grant (JPMJCR23P4), JSPS KAKENHI Grant (23H04830, 22K18265, 823K22379), and MEXT Quantum Leap Flagship Program (MEXT QLEAP) Grant (JPMXS0120330644) to MY.

## Reference

Ahn, W.-Y., Haines, N., & Zhang, L. (2017). Revealing neurocomputational mechanisms of reinforcement learning and decision-making with the hBayesDM package. Computational Psychiatry (Cambridge, Mass.), 1(0), 24–57. 10.1162/CPSY_a_00002

Ang, Y.-S., Gelda, S. E., & Pizzagalli, D. A. (2023). Cognitive effort-based decision-making in major depressive disorder. Psychological Medicine, 53(9), 4228–4235. 10.1017/S0033291722000964

Apps, M. A. J., Grima, L. L., Manohar, S., & Husain, M. (2015). The role of cognitive effort in subjective reward devaluation and risky decision-making. Scientific Reports, 5, 16880. 10.1038/srep16880

Chong, T. T.-J., Apps, M., Giehl, K., Sillence, A., Grima, L. L., & Husain, M. (2017). Neurocomputational mechanisms underlying subjective valuation of effort costs. PLoS Biology, 15(2), e1002598. 10.1371/journal.pbio.1002598

Clay, G., Mlynski, C., Korb, F. M., Goschke, T., & Job, V. (2022). Rewarding cognitive effort increases the intrinsic value of mental labor. Proceedings of the National Academy of Sciences of the United States of America, 119(5), e2111785119. 10.1073/pnas.2111785119

Cutler, J., Contreras-Huerta, L. S., Todorova, B., Nitschke, J., Michalaki, K., Koppel, L., Gkinopoulos, T., Vogel, T. A., Lamm, C., Västfjäll, D., Tsakiris, M., Apps, M. A. J., & Lockwood, P. L. (2025). Psychological interventions that decrease psychological distance or challenge system justification increase motivation to exert effort to mitigate climate change. Communications Psychology, 3(1), 148. 10.1038/s44271-025-00332-4

Escobar, G. G., & Mitchell, S. H. (2025). A systematic review of effort discounting research in humans: Current knowledge, recommendations, and future directions. Judgment and Decision Making, 20(e33), e33. 10.1017/jdm.2025.10009

Gneezy, U., & Rustichini, A. (2000). Pay enough or don’t pay at all. The Quarterly Journal of Economics, 115(3), 791–810. 10.1162/003355300554917

Green, L., Fristoe, N., & Myerson, J. (1994). Temporal discounting and preference reversals in choice between delayed outcomes. Psychonomic Bulletin & Review, 1(3), 383–389. 10.3758/BF03213979

Hauser, T. U., Eldar, E., & Dolan, R. J. (2017). Separate mesocortical and mesolimbic pathways encode effort and reward learning signals. Proceedings of the National Academy of Sciences of the United States of America, 114(35), E7395–E7404. 10.1073/pnas.1705643114

Inzlicht, M., Shenhav, A., & Olivola, C. Y. (2018). The effort paradox: Effort is both costly and valued. Trends in Cognitive Sciences, 22(4), 337–349. 10.1016/j.tics.2018.01.007

Kool, W., & Botvinick, M. (2018). Mental labour. Nature Human Behaviour, 2(12), 899–908. 10.1038/s41562-018-0401-9

Lockwood, P. L., Hamonet, M., Zhang, S. H., Ratnavel, A., Salmony, F. U., Husain, M., & Apps, M. A. J. (2017). Prosocial apathy for helping others when effort is required. Nature Human Behaviour, 1(7), 0131. 10.1038/s41562-017-0131

Lockwood, P. L., Wittmann, M. K., Nili, H., Matsumoto-Ryan, M., Abdurahman, A., Cutler, J., Husain, M., & Apps, M. A. J. (2022). Distinct neural representations for prosocial and self-benefiting effort. Current Biology, 32(19), 4172-4185.e7. 10.1016/j.cub.2022.08.010

Lopez-Gamundi, P., Yao, Y.-W., Chong, T. T.-J., Heekeren, H. R., Mas-Herrero, E., & Marco-Pallarés, J. (2021). The neural basis of effort valuation: A meta-analysis of functional magnetic resonance imaging studies. Neuroscience and Biobehavioral Reviews, 131, 1275–1287. 10.1016/j.neubiorev.2021.10.024

Ma, Q., Meng, L., Wang, L., & Shen, Q. (2014). I endeavor to make it: effort increases valuation of subsequent monetary reward. Behavioural Brain Research, 261, 1–7. 10.1016/j.bbr.2013.11.045

Matthews, J., Pisauro, M. A., Jurgelis, M., Müller, T., Vassena, E., Chong, T. T.-J., & Apps, M. A. J. (2023). Computational mechanisms underlying the dynamics of physical and cognitive fatigue. Cognition, 240, 105603. 10.1016/j.cognition.2023.105603

Moore, M. T., & Fresco, D. M. (2012). Depressive realism: a meta-analytic review. Clinical Psychology Review, 32(6), 496–509. 10.1016/j.cpr.2012.05.004

Müller, T., Klein-Flügge, M. C., Manohar, S. G., Husain, M., & Apps, M. A. J. (2021). Neural and computational mechanisms of momentary fatigue and persistence in effort-based choice. Nature Communications, 12(1), 4593. 10.1038/s41467-021-24927-7

Sayalı, C., & Badre, D. (2021). Neural systems underlying the learning of cognitive effort costs. Cognitive, Affective & Behavioral Neuroscience, 21(4), 698–716. 10.3758/s13415-021-00893-x

Sharot, T. (2011). The optimism bias. Current Biology, 21(23), R941–5. 10.1016/j.cub.2011.10.030

Stan Development Team. (2026). RStan: the R interface to Stan. https://mc-stan.org/

Sutton, R. S., & Barto, A. G. (1998). Reinforcement learning: An introduction. 1(1), 9–11. https://www.cambridge.org/core/journals/robotica/article/robot-learning-edited-by-jonathan-h-connell-and-sridhar-mahadevan-kluwer-boston-19931997-xii240-pp-isbn-0792393651-hardback-21800-guilders-12000-8995/737FD21CA908246DF17779E9C20B6DF6

Taylor, S. E., & Brown, J. D. (1988). Illusion and well-being: A social psychological perspective on mental health. Psychological Bulletin, 103(2), 193–210. 10.1037/0033-2909.103.2.193

Taylor, S. E., & Brown, J. D. (2004). Positive illusions and Weil-being revisited : Separating fact from fiction. Psychological Bulletin. https://taylorlab.psych.ucla.edu/wp-content/uploads/sites/5/2014/10/1994_Positive-Illusions-and-Well-Being-Revisited_Separating-Fact-from-Fiction.pdf

Todorova, B., Zhang, L., Lengersdorff, L., Doell, K. C., Nitschke, J. P., Forbes, P. A. G., Pahl, S., & Lamm, C. (2025). Effort and time costs influence motivational asymmetries in self-benefitting vs pro-environmental decisions. Communications Psychology, 3(1), 166. 10.1038/s44271-025-00347-x

Vassena, E., Silvetti, M., Boehler, C. N., Achten, E., Fias, W., & Verguts, T. (2014). Overlapping neural systems represent cognitive effort and reward anticipation. PloS One, 9(3), e91008. 10.1371/journal.pone.0091008

Vehtari, A., Simpson, D., Gelman, A., Yao, Y., & Gabry, J. (2024). Pareto smoothed importance sampling. In arXiv [stat.CO]. arXiv. 10.48550/arXiv.1507.02646

Westbrook, A., van den Bosch, R., Määttä, J. I., Hofmans, L., Papadopetraki, D., Cools, R., & Frank, M. J. (2020). Dopamine promotes cognitive effort by biasing the benefits versus costs of cognitive work. Science, 367(6484), 1362–1366. 10.1126/SCIENCE.AAZ5891

Westbrook, Andrew, & Braver, T. S. (2015). Cognitive effort: A neuroeconomic approach. Cognitive, Affective & Behavioral Neuroscience, 15(2), 395–415. 10.3758/s13415-015-0334-y

Yantis, S., Schwarzbach, J., Serences, J. T., Carlson, R. L., Steinmetz, M. A., Pekar, J. J., & Courtney, S. M. (2002). Transient neural activity in human parietal cortex during spatial attention shifts. Nature Neuroscience, 5(10), 995–1002. 10.1038/nn921

